# Evaluation of the effects of library preparation procedure and sample characteristics on the accuracy of metagenomic profiles

**DOI:** 10.1101/2021.04.12.439578

**Authors:** Christopher A Gaulke, Emily R Schmeltzer, Mark Dasenko, Brett M. Tyler, Rebecca Vega Thurber, Thomas J Sharpton

## Abstract

Shotgun metagenomic sequencing has transformed our understanding of microbial community ecology. However, preparing metagenomic libraries for high-throughput DNA sequencing remains a costly, labor-intensive, and time-consuming procedure, which in turn limits the utility of metagenomes. Several library preparation procedures have recently been developed to offset these costs, but it is unclear how these newer procedures compare to current standards in the field. In particular, it is not clear if all such procedures perform equally well across different types of microbial communities, or if features of the biological samples being processed (e.g., DNA amount) impact the accuracy of the approach. To address these questions, we assessed how five different shotgun DNA sequence library preparation methods, including the commonly used Nextera^®^ Flex kit, perform when applied to metagenomic DNA. We measured each method’s ability to produce metagenomic data that accurately represents the underlying taxonomic and genetic diversity of the community. We performed these analyses across a range of microbial community types (e.g., soil, coral-associated, mouse-gut-associated) and input DNA amounts. We find that the type of community and amount of input DNA influence each method’s performance, indicating that careful consideration may be needed when selecting between methods, especially for low complexity communities. However, cost-effective preparation methods we assessed are generally comparable to the current gold standard Nextera^®^ DNA Flex kit for high-complexity communities. Overall, the results from this analysis will help expand and even facilitate access to metagenomic approaches in future studies.

**IMPORTANCE:** Metagenomic library preparation methods and sequencing technologies continue to advance rapidly, allowing researchers to characterize microbial communities in previously underexplored environmental samples and systems. However, widely-accepted standardized library preparation methods can be cost-prohibitive. Newly available approaches may be less expensive, but their efficacy in comparison to standardized methods remains unknown. In this study, we compared five different metagenomic library preparation methods. We evaluated each method across a range of microbial communities varying in complexity and quantity of input DNA. Our findings demonstrate the importance of considering sample properties, including community type, composition, and DNA amount, when choosing the most appropriate metagenomic library preparation method.

## INTRODUCTION

Recent advancements in high-throughput sequencing have revolutionized genomic discovery and unlocked new insights regarding the diversity and function of microbial communities (1–4). For example, shotgun metagenomic sequencing has clarified how the functional capacity of the gut microbiome links to human health (5–8), improved the efficacy of antibiotic resistance gene discovery (9–12), identified beneficial soil microbes for agricultural use (13–15), and uncovered novel, medically relevant biosynthetic gene clusters in marine microbes (16–18). However, while metagenomes offer rich opportunity to transform discovery, the financial cost of producing metagenomic data limits their application. Because much of this cost is associated with the preparation of metagenomic DNA for high-throughput sequencing, there is hope that emergent economical products and procedures can expand the scope of metagenomic investigations.

Illumina’s Nextera^®^ XT and DNA Flex kits (the latter now known as the “Illumina^®^ DNA Prep”) have been the most widely used methods for preparing metagenomic libraries and have effectively served as industry standard approaches. Indeed, Illumina DNA sequencing platforms remain the most widely utilized for generating genomic and metagenomic data, and their library preparation kits are accordingly used to prepare samples for sequencing. Due to their frequent use, these kits are subject to extensive evaluation and refinement. For example, Illumina recently released an updated version of their “gold standard” Nextera^®^ XT kit, which was rebranded as the Nextera^®^ DNA Flex (and now Illumina^®^ DNA Prep). This new kit allows greater flexibility across a wider range of genomes, from small genomes (microbial and amplicons) to more complex genomes found in eukaryotic and human systems. The Flex kit also resolved sequencing biases identified in the Nextera^®^ XT kit that occur in genomic regions with extreme GC-content (19). These features of the Nextera^®^ DNA Flex kit have contributed to its broad adoption in metagenomic investigations.

One downside to the Nextera^®^ DNA Flex kit is its relatively high price, which presently costs roughly $46 per sample. While this cost may be reasonable considering the demand for the product and its observed efficacy, it is high enough that it limits the scale of many metagenomic investigations. For example, studies performing high-throughput analyses on hundreds or thousands of samples may be forced to utilize non-metagenomic approaches (e.g., 16S rRNA gene sequencing) due to this library preparation expense. In the effort to circumvent this challenge, several alternative and competitive genomic library preparation methods have recently been developed and applied to metagenomic investigations. These approaches fall into two categories: methods that increase the economy of the Illumina Nextera^®^ by modifying various aspects of the manufacturing protocols (e.g. Baym et al. 2015 (20), and those that use entirely different technologies (e.g. seqWell plexWell™ 96). These approaches hold great promise to improve the throughput of metagenomic investigations by reducing library preparation costs. For example, the recent method known as ‘Hackflex’ achieves an eleven-fold decrease in per sample reagent costs compared to the Illumina kit protocols (21).

Although several alternative library preparation approaches have been assessed from the perspective of whole genome sequencing, very little is known about their accuracy and precision when applied to metagenomic investigations. It is crucial that the performance of novel library preparation procedures be specifically assessed in diverse metagenomic communities as different community types provide the unique sequencing challenges not common to traditional whole genome sequencing. For example, metagenomic communities vary in complexity, with some communities having few distinct taxa (e.g., insect gut) while others are very highly diverse (e.g., soil). Library preparation procedures may vary in their ability to unbiasedly sample DNA across the different genomes present in the community. Biological samples vary in their biomass, which affects the amount of whole community DNA that is subject to the library preparation approach. The sensitivity of these approaches to the amount of input DNA may hence impact study outcomes (25–28).

To advance the utility of low cost metagenomic library preparation methods, we quantified the performance of five recently developed approaches. Our investigation assessed how different features of metagenomic samples, including community complexity, and biomass impacts the performance of these procedures. In particular, we compared the Illumina Nextera^®^ DNA Flex, a modified DNA Flex protocol (20), QIAGEN^®^ QIAseq FX DNA, Perkin Elmer NEXTFLEX^®^ Rapid DNA-Seq 2.0, and seqWell plexWell™ 96 library preparation methods using community acquired DNA obtained from three different types of microbial communities: low complexity communities (as represented by *Acropora hyacinthus* microbiomes), moderately complex communities (as represented by *Mus musculus* mouse fecal microbiomes), and a highly complex community (as represented by a soil microbiome). We also evaluated how each approach performs on a commercially available mock community comprised of ten microbial species. Our analysis clarifies the performance of these approaches across these different sample conditions and the results will assist investigators in identifying appropriate approaches for their metagenomic investigations.

## RESULTS

### Library preparation procedure, community type, and input concentration influence metagenomic library characteristics

To determine how metagenomic library characteristics (e.g., insert size, millions of sequences generated) varied across different metagenomic library preparations, we regressed each library characteristic on community type, library preparation, and input DNA concentration (Table 1). We found that these predictor variables statistically affected the following characteristics: median fragment size (F_(31,48)_ = 29.94, R^2^ = 0.90, *P* < 2.2×10^−16^), library concentration (F_(31,48)_ = 12.44, R^2^ = 0.82, *P* = 3.51×10^−14^), library molarity (F_(31,48)_ =11.75, R^2^ 0.81, *P* = 1.09×10^−13^), sequence read length (F_(39,60)_= 56.81, R^2^ = 0.96, *p* < 2.2×10^−16^), number of reads generated (F_(39,60)_ = 11.69, R^2^ = 0.81, *P* < 2.2×10^−16^), read GC content (F_(39,60)_ = 2285, R^2^ = 0.99, P < 2.2×10^−16^), duplication rate (F_(8,91)_ = 2.494, R^2^ = 0.11, *P* = 0.02), and percentage of reads filtered (F_(39,60)_ = 1716, R^2^ = 0.99, *P* < 2.2×10^−16^). Sequence duplication rate was only sensitive to community type (F_(3,91)_ = 5.93, *P* = 9.0×10^−04^). Specifically, communities with low microbial diversity such as the coral (t = 2.885, *P* = 4.88×10-03) and mock communities (t = 2.16, *P* = 0.03) had elevated duplication rates. All other library characteristics were sensitive to interactions between community type, library preparation, and input DNA concentration, and many characteristics were also impacted by the independent effects of these variables. For example, library preparation method (F_(3,48)_ = 148.85, *P* = 2.61×10^−24^) and community type (F_(3,48)_ = 11.83, *P* = 6.39×10^−06^) affected median fragment size independent of the interaction between these parameters and DNA input (F_(9,48)_ = 5.93, *P* = 1.56×10^−05^). Library concentration was impacted by community type (F_(3,48)_ = 5.72, *P* = 1.98^−03^), library preparation (F_(3,48)_ = 20.75, *P* = 9.22×10^−09^), input DNA concentration (F_(1,48)_ = 186.86, *P* < 2.2×10^−16^) and their interaction (F_(9,48)_ = 6.09, *P* = 1.16×10^−05^). Library molarity was similarly impacted by community type (F_(3,48)_ = 5.63, *P* = 2.18×10^−03^), library preparation (F_(3,48)_ = 35.69, *P* = 2.82×10^−12^), input DNA concentration (F_(1,48)_ = 126.21, *P* = 4.89×10^−15^) and their interaction (F_(9,48)_ = 4.40, *P* = 3.16×10^−04^).

**Table 1.**
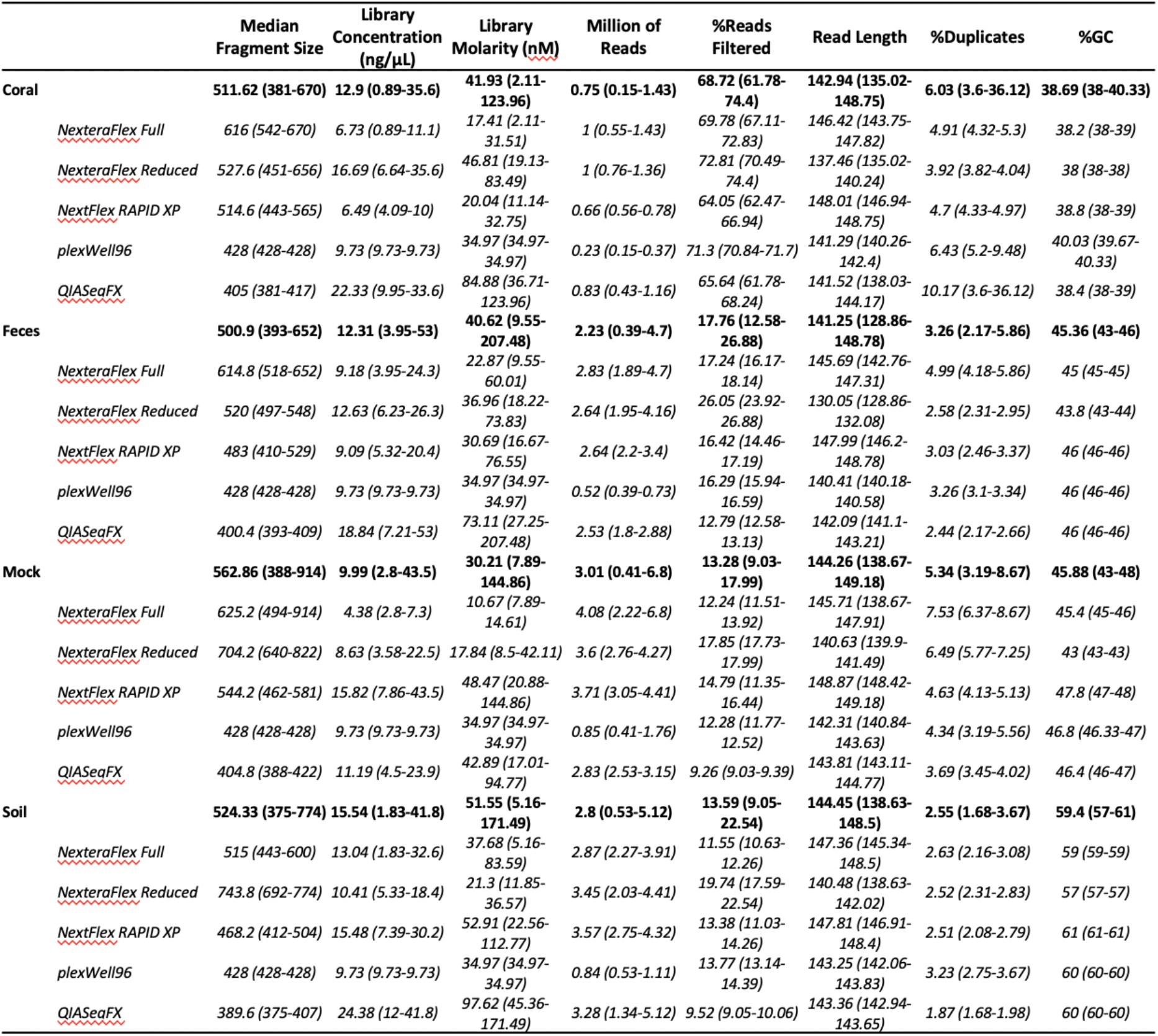
Library preparation summary statistics. Each column show means and range of NGS library parameters in coral, feces, mock, and soil sample types. Parameters that varied significantly (p < 0.05) by community type(*), input concentration (†), and library preparation(‡)

The number of sequences generated was sensitive to preparation procedure (F_(4,60)_ = 41.68, *P* = 1.10×10^−16^), community type (F_(3,60)_ = 67.67, *P* = 3.09×10^−19^), and DNA input concentration (F(1,60) = 5.62, *P* = 2.09×10^−02^) as well as the interaction between these variables (F_(12,60)_ = 2.25, *P* = 2.00×10^−02^). The number of sequence reads that were quality filtered and derived from the host genome was also significantly affected by this interaction (F_(12,60)_ = 5.16, *P* = 7.44×10^−06^). Metagenome libraries constructed for coral communities had increased levels of quality filtering (t = 62.11, *P* = 3.66×10^−56^) while filtering in soil (t = −6.65, *P* = 9.84×10^−09^) and mock (t = −6.56, *P* = 1.39^−08^) communities was decreased when compared to fecal samples. Read filtering was also consistently increased in samples prepared with the Nextera^®^ Flex Reduced method (t = −12.38, *P* = 3.59^−18^) and reduced in the samples prepared using the QIASeqFX procedure (t = −4.89, *P* = 7.84^−06^).

Metagenomic read characteristics were also significantly impacted by the examined variables. For example, the GC content of reads was significantly dependent on community type (F_(3,60)_ = 28,941.39, *P* = 9.38×10^−95^). GC content was also affected by library preparation method (F_(4,60)_ = 442.80, *P* = 8.71×10^−44^); a significant effect of the interaction between library preparation, community type, and input DNA concentration (F_(12,60)_ = 2.68, *P* = 5.98×10^−03^) was also observed for GC content. Finally, average read length after quality filtering varied by both community type (F_(3,60)_ = 60.78, *P* = 3.54×10^−18^) and preparation procedure (F_(4,60)_ = 404.25, *P* = 1.21×10^−42^). Collectively, these findings indicate that metagenomic library preparation procedures yield distinct library characteristics for different community types.

### Different library preparation methods result in similar taxonomic profiles of a standardized mock community

While library preparation procedures varied in the resulting metagenomic library and sequence characteristics, it is unclear if this variation results in different downstream assessment of community composition. To address this question, we quantified how each library preparation method predicted the taxonomic composition of a defined mock community. We compared the taxonomic composition generated by each library preparation to the ZymoBIOMICS™ Microbial Community standard’s defined taxonomic composition of the mock community. Strong correlations (ρ = 0.93 - 0.97, *P* = 1.29×10^−6^ - *P* < 2.2×10^−16^, fdr < 1.0×10^−5^) were observed between the MetaPhlAn2 inferred taxonomic abundances and theoretical taxonomic abundances of taxa present in the mock community (Figure 1). To confirm that this was not due to bias in the MetaPhlAn2 database, we also compared the inferred taxonomic abundances using Kraken2 and observed similar taxonomic associations (ρ = 0.93 - 0.96, all *P* < 2.2×10^−16^, fdr < 2.2×10^−16^) and abundance profiles (Supp. Fig. 1).

**Figure 1.**
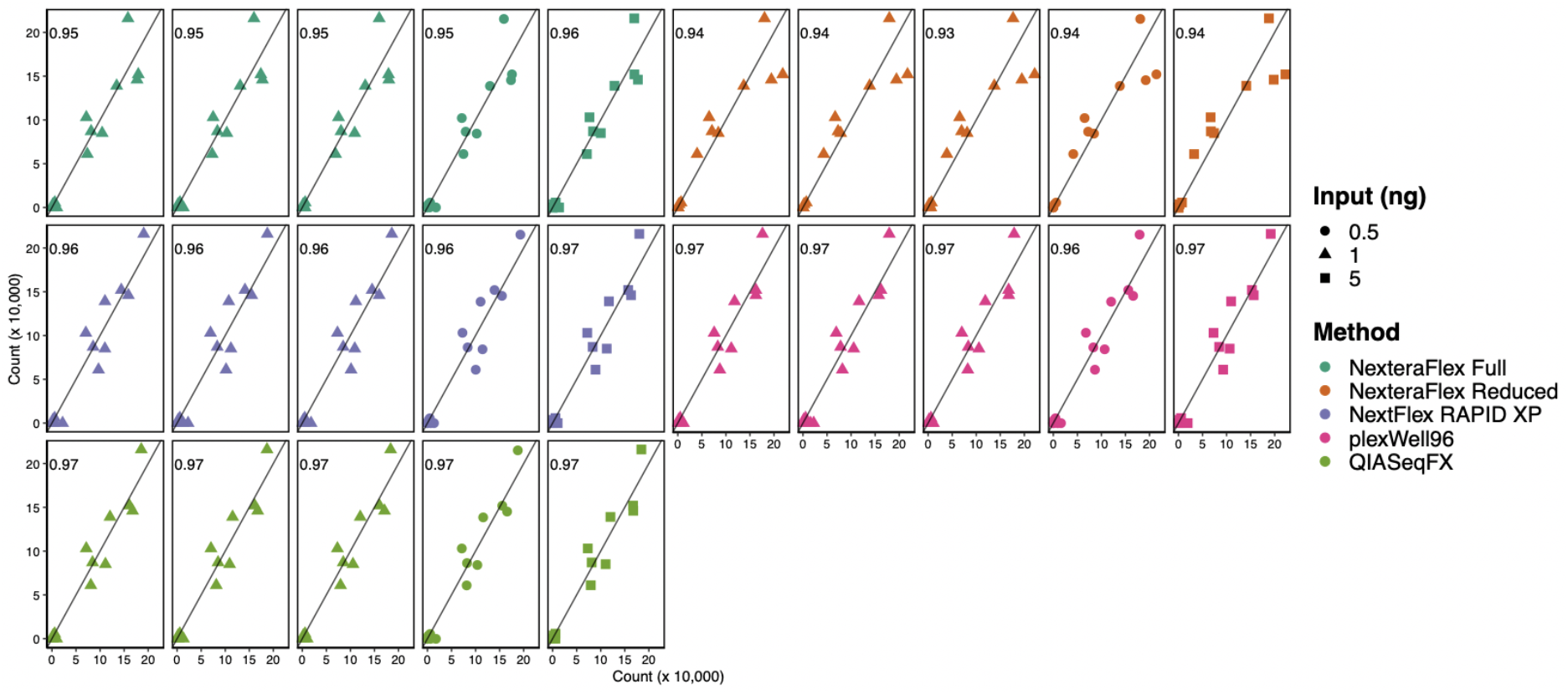
Metagenomic library preparation methods accurately predict taxonomic composition of a simplified mock community. Scatter plots of the observed and theoretical taxonomic compositions of a mock community. Libraries preparation methodology is indicated by point color and input concentration is denoted with point shape. Values in each panel represent the Pearson correlation coefficient.

MetaPhlAn2 results showed that the Nextera^®^ Flex Full and Nextera^®^ Flex Reduced methods, which are widely used in metagenomic studies, had the lowest correlations (ρ = 0.93 – 0.96, *P* = 1.29×10^−6^ - 2.66×10^−8^) with the theoretical composition of the mock community. The lower correlations produced by these two library preparation procedures are driven by an underestimation of *Lactobacillus fermentum* and an overestimation of the abundance of *Staphylococcus aureus* and *Enterococcus faecalis*. The strongest correlations were observed in the QIASeqFX (ρ = 0.97, *P* = 1.21×10^−8^ - 5.79×10^−9^), plexWell96 (ρ = 0.96 - 0.97, *P* = 2.58×10^−8^ - 1.24×10^−8^), and NEXTFLEX Rapid (ρ = 0.96 - 0.97, *P* = 1.02×10^−7^ - 2.38×10^−8^) methods (Figure 1). Together these data indicate that library preparation methods subtly influence some taxonomic estimates but that all methods examined overall performed well at recapitulating simple, defined microbial communities.

### Community taxonomic profiles are significantly impacted by library preparation procedure and input concentration

Next, we quantified the impact of library preparation methods and input concentrations on the taxonomic profiles of soil, coral, mock, and fecal metagenomes. Library preparation procedure significantly associated with resulting taxonomic microbiome profiles as measured by PERMANOVA in coral (R^2^ = 0.70, *P =* 2.00×10^−4^), feces (R^2^ = 0.85, *P =* 2.00×10^−4^), mock (R^2^ = 0.72, *P =* 4.00×10^−4^), and soil (R^2^ = 0.72, *P =* 4.00×10^−4^) (Figure 2A) communities. In the fecal, soil, and mock communities, no association was found between input concentration and microbiome beta-diversity. However, in coral communities we identified a significant interaction effect between input concentration and metagenomic preparation method (R^2^ = 0.13, *P* = 3.0×10^−2^). The taxonomic abundance profiles of each library were highly correlated (ρ = 0.63 – 1, fdr < 2.2×10^−16^) across library preparation methods (Figure 2B) suggesting that preparation methodology associates with distinct community profiles.

**Figure 2.**
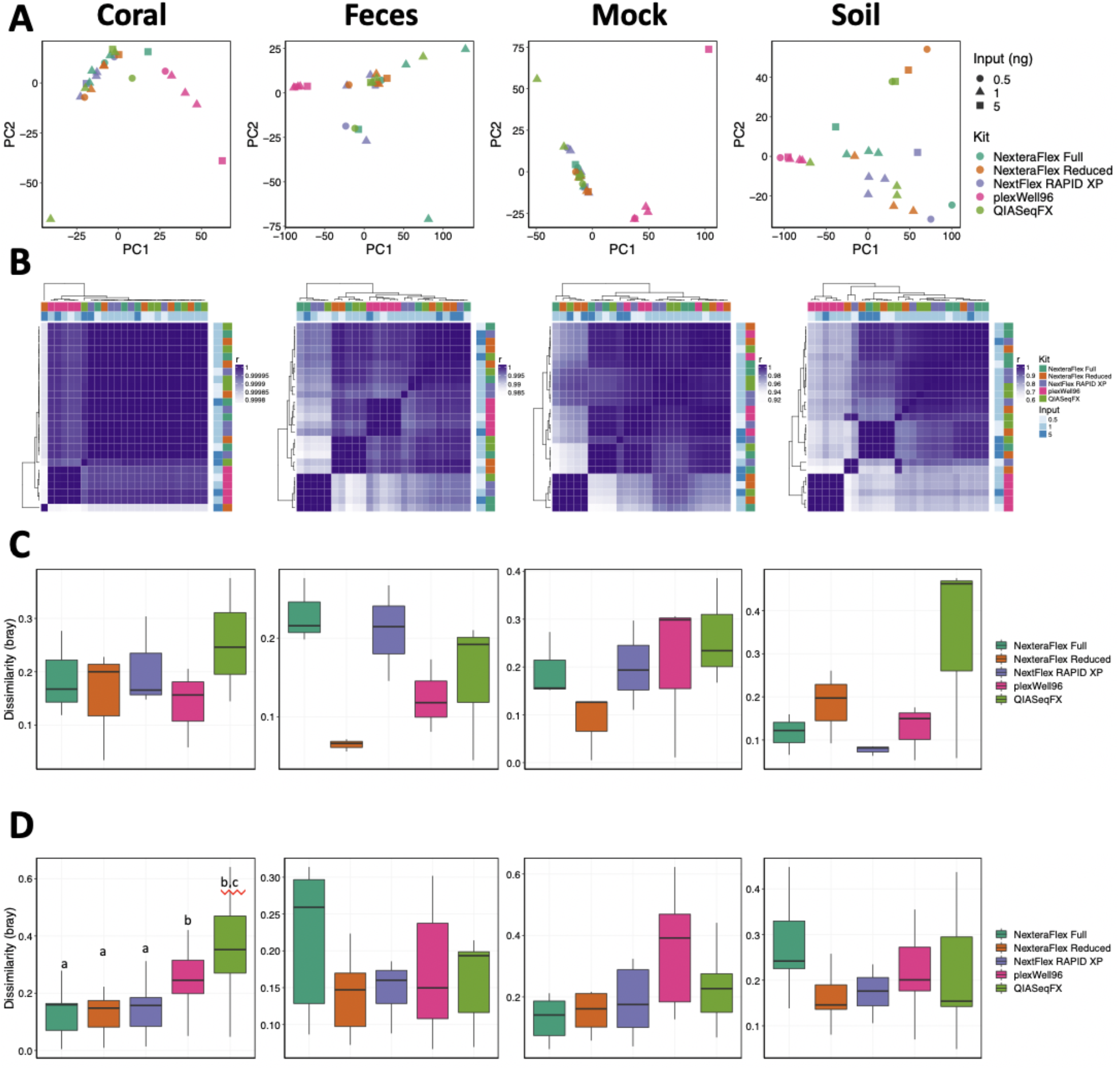
Microbial taxonomic diversity varies by library preparation method and input concentration. **A)** PCA ordinations of microbial taxonomic diversity for each sample type. **B)** Correlation heat maps of taxonomic abundances generated from each library type. Row and column side plots indicated the library preparation methodology and the input concentration. Boxplots of dissimilarities (Bray Curtis) between **C)** technical replicates (lng inputs), and **D)** different input concentrations. Letters indicate significant differences, *P* < 0.05.

To quantify how variable individual replicates were across different library preparation methods, we examined the dissimilarity in taxonomic beta-diversity within each library preparation method. Methods that yield low intra-concentration (i.e., 1 ng/μL input) dissimilarities indicate high levels of reproducibility, while methods with low inter-concentration dissimilarities (i.e., all concentrations) would indicate that the taxonomic profiles generated using this method are robust to variation in library input concentration. We found that variation in intra-concentration taxonomic dissimilarity was low and significant differences were not observed across library preparation methodologies (Figure 2C). Inter-concentration dissimilarity was also low in fecal, mock, and soil samples but varied significantly across preparation methods in coral (H = 12.28, *P* = 0.02) potentially due to elevated dissimilarity in the QIASeqFX libraries. Together these data suggest that the taxonomic profiles generated using the methods under investigation are similarly reproducible, but the robustness varied across methods.

### Community gene abundance profiles are sensitive to library preparation procedures and input DNA concentration

Metagenomic investigations frequently seek to define the genetic diversity of microbial communities. Using the number of distinct gene families observed in the data (i.e., gene family richness) as well as the functional composition of the community (i.e., gene family beta-diversity), we measured how different library preparation procedures affected determination of a community’s functional capacity. Gene family richness varied by library preparation method and input concentration in coral (F_(9,15)_ = 31.90, R^2^ = 0.92, *P* = 3.47 ×10^−08^), feces (F_(9,15)_ = 9.13, R^2^ = 0.75, *P* = 1.20 ×10^−04^), and mock (F_(9,15)_ = 8.78, R^2^ = 0.74, *P* = 1.51 ×10^−04^) communities, while soil richness (F_(9,15)_ = 1.41, R^2^ = 0.13, *P* =0.27) was less sensitive to these effects (Figure 3A). We also observed a significant interaction between library preparation procedure and DNA input on the predicted functional profiles of coral (F_(4,15)_ = 8.00, *P* = 1.16 x 10^−03^), feces (F_(4,15)_ = 5.64, *P* = 5.60 x 10^−03^), and mock microbiomes (F_(4,15)_ = 7.78, *P* = 1.33 x 10^−03^). However, these associations were not always consistent across different library preparation procedures. For example, gene richness was elevated in coral samples prepared with the plexWell™96 method (t = 5.53, *P* = 5.77×10^−05^), while a contrasting pattern was observed in both fecal (t = −6.38, *P* = 1.23 ×10^−05^) and soil (t = −1.958, *P* = 6.91 ×10^−02^) samples, and no difference was identified in mock community samples (t = −0.58, *P* = 0.57). Similar effects of library preparation method and input concentration were observed on Shannon entropy (Figure 3B). Specifically, significant effects were observed for coral (F_(9,15)_ = 11.09, R^2^ = 0.79, *P* = 3.75 ×10^−05^), feces (F_(9,15)_ = 62.29, R^2^ = 0.96, *P* = 1.89 ×10^−10^), mock (F_(9,15)_ = 11.04, R^2^ = 0.79, *P* = 3.85 ×10^−05^), and soil (F_(9,15)_ = 5.06, R^2^ = 0.60, *P* = 2.95 ×10^−03^).

**Figure 3.**
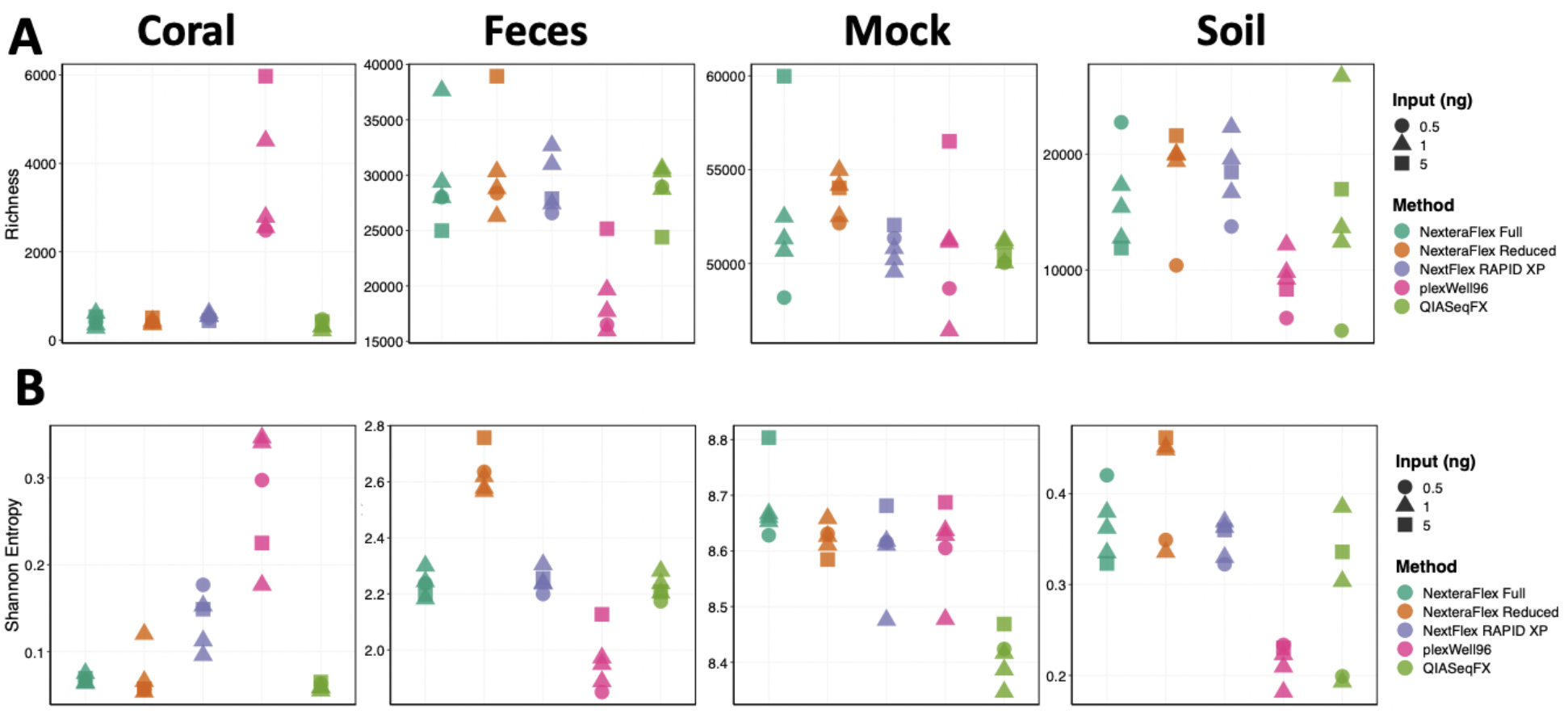
Metagenomic diversity varies by library preparation method and input concentration. **A)** Gene and **B)** Shannon entropy plots for each sample type.

As measured by PERMANOVA, we found that gene family beta-diversity (Bray Curtis) was significantly associated with library preparation method for coral (F_(4,15)_ = 3.62, R^2^ = 0.44, *P* = 2.00 ×10^−04^), fecal (F_(4,15)_ = 7.82, R^2^ = 0.60, *P* = 2.00 ×10^−04^), mock (F_(4,15)_ = 3.36, R^2^ = 0.41, *P* = 2.00 ×10^−04^), and soil (F_(4,15)_ = 3.39, R^2^ = 0.40, *P* = 2.00 ×10^−04^) communities (Figure 4A). However, the association between library preparation procedure and gene family beta-diversity is muted in comparison to taxonomic beta-diversity (Figure 2A), possibly as a result of the increased overall similarity in gene family abundances across samples of the same community type (ρ = 0.99 – 1.00, *P* < 2.2×10^−16^; Figure 4B). Despite this increased similarity between library preparation methods for gene family abundances, the beta-diversity of technical replicates did vary across library preparation methods for coral (H = 11.43, *P* = 2.21 x 10^−02^), fecal (H = 9.03, *P* = 6.02 x 10^−02^), mock (H = 12.1, *P* = 1.66 x 10^−02^), and soil (H = 11.5, *P* = 2.15 x 10^−02^) samples.

**Figure 4.**
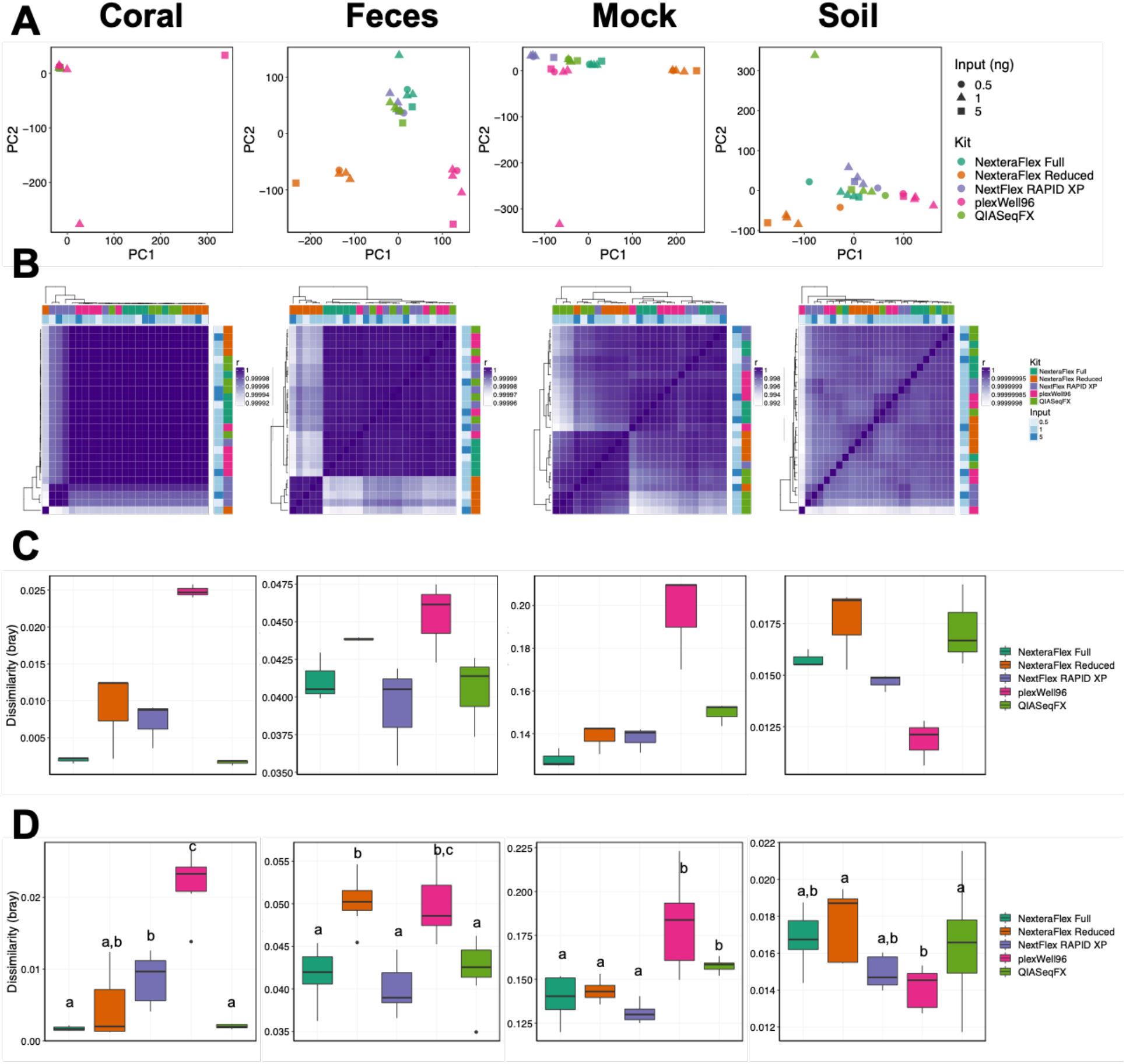
Microbial functional diversity varies by library preparation method and input concentration. **A)** PCA ordinations of gene family beta-diversity for each sample type. **B)** Correlation heat maps of gene family abundances generated from each library type. Row and column side plots indicated the library preparation methodology and the input concentration. Boxplots of dissimilarities (Bray Curtis) between **C)** technical replicates (1ng inputs), and **D)** different input concentrations. Letters indicate significant differences, *P* < 0.05.

However, significant differences were not detected between individual library preparation methods (Figure 4C). The robustness of metagenome beta-diversity to input concentration differed across library preparation methods for coral (H = 24.33, *P* = 6.86 x 10^−05^), fecal (H = 24.53, *P* = 6.26 x 10^−05^), mock (H = 25.65, *P* = 3.72 x 10^−05^), and soil (H = 13.83, *P* = 7.85 x 10^−03^) samples (Figure 4D). Notably, the plexWell™96 method had elevated variability compared to the other library prep methods in coral, fecal, and mock community samples. In soil, these patterns were mitigated (Figure 4D). Overall, these data demonstrate that, similar to the taxonomic profiles, library preparation methods affect gene diversity profile predictions.

## DISCUSSION

### Taxonomic and functional profiles predictions are similar across methodology

Although the Nextera^®^ kits are widely used and considered the ‘gold standard’ for metagenomic sample preparation, their cost can limit researchers from conducting expansive project aims. As applications for metagenomic sequencing continue to increase, researchers are left with the difficult task of balancing the need for high-quality data with the cost of its generation. The development of new protocols that modify the standard Nextera^®^ kit protocol as well as several new economical library preparation kits have the potential to dramatically alter the field by expanding the accessibility of shotgun metagenomics. However, the quality of libraries prepared using more economical methods varies substantially (29). While prior studies have demonstrated that different library preparation procedures can affect metagenome characteristics (30–33), these studies did not evaluate contemporary procedures nor did they consider the sensitivity of the approaches to different metagenome sample types. Here we demonstrate that library quality as well as taxonomic and functional profiles vary as a function of environmental community type and biomass. Our findings suggest that while researchers need to be aware of differences between kits, overall taxonomic and functional profiles produced by these kits are similar.

Several investigations have identified key differences in library characteristics across metagenomic library preparation procedures, often by incorporating multiple study designs. These variations can result in substantial changes in the quality of the metagenomic library and are important considerations in preparation method selection. For example, Baym et al. demonstrated that a custom Nextera^®^ XT protocol yielded a substantially reduced insert fragment length (20). Smaller fragments have higher proportions of adapter contamination in reads, while fragments that are too large may be preferentially lost during the Illumina cluster generation process (34). We observed significant effects of community type, library preparation procedure, and input DNA concentration on fragment size. In our hands, the Nextera^®^ Flex protocols generated the largest library insert sizes, while the QIAseq FX and plexWell™96 procedures consistently produced the smallest. However, the Nextera^®^ procedures also produced libraries with the lowest average GC content when compared to the other procedures examined. This reduced representation of GC could impact the representation of genes with high GC content and skew both taxonomic and functional profiles (29, 35). The interacting effects of library preparation procedure, community type, and DNA input on GC content further indicates that specific library preparation procedures may have distinct insertion site biases.

Comparing library characteristics across environmental sample types, samples with low relative diversity (i.e., coral) had both a higher percentage of duplicate reads and a high number of reads filtered and removed from the resulting libraries regardless of input concentration. This high level of filtering is likely due to the extreme levels of host DNA contaminants relative to the other sample types, and additionally may point to the larger issue of host sequence contamination, regardless of library preparation method, in similar research studies. However, coral samples also had similar levels of precision across kit types, with the exception of lower precision with QIASeq FX, demonstrating that different kit types may still be viable options in other low complexity study systems.

Samples with moderate (fecal) and high (soil) microbial diversity had much lower respective average percentage of sequencing reads filtered than coral samples, but with a higher average GC content across libraries than coral samples. Fecal sample libraries had the highest variability in precision between kit types, with the exception of Nextera^®^ Flex Reduced, likely due to the more complex community composition. However, our study also had a relatively low average sequencing depth across all samples due to sample number and financial constraints; the high intra-community variability in precision that we observed may be resolved with a higher sequencing coverage (37). It is also possible that longer insert fragment sizes introduce greater variability due to lower base quality in produced community composition regardless of sample type (38), though this was not the case for coral, soil, or mock libraries with the longest fragment sizes.

In successfully recapitulating the taxonomic profiles of a mock microbial community, all library preparation methods performed similarly overall, however, variation in taxonomic profiles for the environmental sample types showed subtle differences between methods. While higher levels of intra-community variation per method could again be due to low sequencing depth, our results of higher variation for the coral sample types with lower relative diversity are consistent with previous findings that library coverage is increased for highly complex microbial communities. Furthermore, while it may appear that all preparation methods perform poorly in both taxonomic and functional resolution for low (coral) and high (soil) diversity sample types, it must be noted that these profiles may only be as complete as the reference databases used for assignment, and it is well known that these databases are preferentially curated with human microbiome sequences and studies in mind (39).

### Financial and opportunity costs of metagenomic preparation methods differ

Decreasing the costs of kits and reagents associated with library preparation improves access to metagenomic approaches. The Nextera^®^ DNA Flex Full preparation actualized cost remains the most expensive of the five methods tested, with the NEXTFLEX Rapid XP and QIASeq FX in the median relative expense range, and Nextera^®^ Flex Reduced prep and plexWell™96 as the most economical choices for metagenomic library generation. However, due to the noted effect of specificity of environmental sample type on performance of preparation method, neither the most economical choice nor the most expensive may necessarily suit every study or generate the highest quality libraries. Due to the effects of preparation procedure, community type, and DNA input on fragment size and both taxonomic and functional profiles of metagenomic samples, comparing communities across multiple study designs may require additional covariates in statistical design. For future studies we recommend incorporating library preparation technique as a potential covariate in statistical design to account for these known differences and potential biases.

### Conclusion

Collectively, these findings demonstrate that no single metagenomic library preparation approach performed the best across all community types and conditions evaluated. Rather, the performance of approaches varied as function of sample and the amount of input DNA. Consequently, researchers should consider these variables when selecting library preparation approaches, especially when attempting to optimize data quality, accuracy, and precision. To aid in this effort, we provide Figure 5 as a reference guide to aid in choosing preparation methods with cost and performance in mind. Our hope is that this information helps improve the accessibility and utility of metagenomic investigations. Further study is needed to determine what community properties (e.g., GC content, taxonomic diversity, etc.) dictate these differences in library procedure performance in order to generate more generalizable guidance of procedure selection. That said, our results show that the different approaches generally produced relatively consistent taxonomic and gene family diversity profiles, which indicates that selecting approaches based on cost and ease of implementation may be appropriate for some studies (namely those in which the loss of accuracy and precision is tolerable). However, we recommend careful consideration of community type and the amount of input DNA when selecting a metagenomic library preparation procedure to ensure optimal performance.

**Figure 5.**
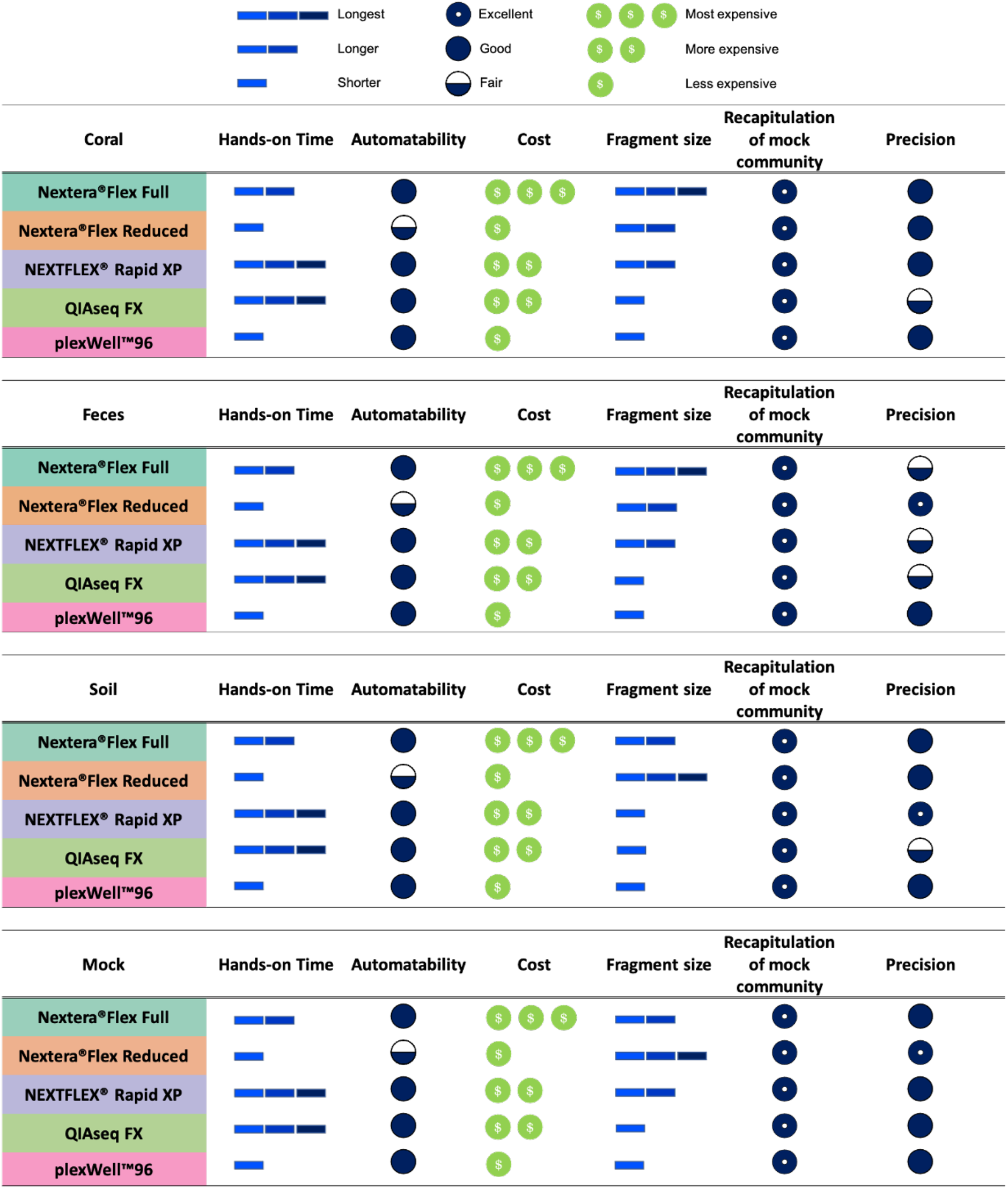
Library preparation summary and cost metrics reference guide. Hands-on time refers to active time necessary for essential benchwork tasks. Fragment size categories are relative to other kit fragment sizes for each sample type. Recapitulation of mock community refers to correlation coefficient of given mock community and community produced by each kit. Precision refers to the level of variability in taxonomic community composition between 1.0ng DNA input technical replicates for each sample type.

## MATERIALS AND METHODS

### Genomic DNA extraction

Prior to metagenomic library construction, genomic DNA was extracted from environmental samples originating from: soil from the North American Project to Evaluate Soil Health Measurements (40), coral (*Acropora hyacinthus*), and mammalian feces (*Mus musculus*; C57BL/6) using methods outlined below. In addition to environmental samples, we used the ZymoBIOMICS™ Microbial Community DNA Standard to more efficiently assess bias associated with library preparation methods on a standard mock community.

For coral slurry, coral nubbins preserved in RNA/DNA Shield (ZymoBIOMICS™) were vortexed in 15ml tubes with a combination of ceramic and garnet bead lysing matrix at ~2500 RPM for 25 minutes. DNA was extracted from 300μl of the resulting coral slurry using ZymoBIOMICS™ DNA/RNA Miniprep (Zymo Research Corp., Irvine, CA, USA) following an additional 2-step enzyme incubation to increase recovery of bacterial DNA: 1) addition of 30μl chicken egg-white lysozyme (10mg/ml, Novagen^®^), 1.8μL mutanolysin (50KU/ml from *Streptomyces globisporus* ATCC 21553, Sigma-Aldrich), 1.8μL lysostaphin (4KU/ml from *Staphylococcus staphylolyticus,* Sigma-Aldrich) and incubation at 37°C for 1 hr and 2) 1 hour incubation at 50°C following addition of 15μl Proteinase K (20 mg/ml, Thermo Scientific™) and 30μl Proteinase K digestion buffer (0.1M NaCl, 10mM Tris pH 9.0, 1mM EDTA, 0.5% SDS, nuclease-free water). Following digestion, one volume of kit-specific DNA/RNA Lysis Buffer was added in order to proceed with the manufacturer’s recommended extraction protocol.

For soil, the sample was taken on 2/27/2019 at the Virginia Tech Eastern Shore Agricultural Research and Extension Center. Samples were collected as 12 composite knife slices of soil to a depth of 15 cm, and each of the 12 slices were passed through an 8mm filter. Detailed sampling methods can be reviewed in Norris et al. 2020 (40). Following collection, 0.25g aliquots of soil were stored at −80°C after overnight shipment from the collection site. Soil aliquots were then extracted following the Earth Microbiome Protocol (41) using a KingFisher™ Flex (Thermo Fisher^®^).

For mouse feces, DNA was isolated from a single fecal pellet using the DNeasy PowerSoil isolation kit (QIAGEN^®^) following the manufacturer’s instructions. An additional 10-minute incubation at 65°C directly before bead beating was added to enhance microbial cell lysis. The samples were then homogenized using a Vortex-Genie 2 and vortex adapter (QIAGEN^®^) at the highest setting for 10 minutes.

### Metagenomic library preparation and sequencing

Environmental DNA samples were prepared for metagenomic sequencing following manufacturers’ protocol using four commercially available kits: 1) Illumina Nextera^®^ DNA Flex Library Kit, 2) QIAGEN^®^ QIAseq FX DNA Library Kit, 3) Perkin Elmer NEXTFLEX^®^ Rapid DNA-Seq Kit 2.0, and 4) seqWell plexWell™ 96. In addition, we included a fifth preparation method using the modified “reduced” protocol established by Baym et al. to increase the number of libraries the Nextera^®^ DNA Flex could generate (20). Genomic DNA was quantified using a Qubit 1X dsDNA HS Assay Kit for soil, fecal, and coral communities. Mock community DNA concentration was not quantified as ZymoBIOMICS™ manufacturer information provided a known concentration of 100ng/μl. Following quantification, all samples prepared using Nextera^®^Flex Full, QIASeq FX, and NEXTFLEX^®^ Rapid XP were diluted with water to 0.2ng/μl. To determine how DNA input affected library generation, each standardized DNA concentration was then added to obtain respective 0.5ng, 1.0ng, and 5.0ng inputs. Samples prepared using plexWell™96 were diluted to 0.25ng/μl with water and appropriate additive volumes made to obtain 0.5ng and 1.0ng input concentrations. For samples with 5.0ng inputs, samples were diluted to 1.25ng/μl and then 4μl of sample was used to obtain 5.0ng input concentration. For the Nextera^®^Flex Reduced reaction, all samples were diluted to 5.0ng/μl with nuclease-free water. A 1μl aliqout of this dilution was used for 5.0ng input libraries. For 0.5ng and 1.0ng input libraries, sufficient water was added to the 1μl dilution to bring respective concentrations to 0.5ng/μl and 1.0ng/μl.

Library insert size was assessed for the Nextera^®^ DNA Flex (full and reduced) and plexWell™96 methods using the Agilent TapeStation 4200 high sensitivity D5000 DNA ScreenTape. Insert size for the QIAseq FX and NEXTFLEX^®^ Rapid XP methods was quantified using the Agilent Bioanalyzer 2100 high-sensitivity DNA chip as these libraries are more prone to have adapter dimers which are poorly resolved using the TapeStation. Library concentration was assessed with the Qubit 1X high-sensitivity dsDNA quantification kit (ThermoFisher). Resulting libraries were diluted, pooled, and sequenced for paired-end reads of 150bp on Illumina HiSeq3000.

### Microbial community gene family abundance and taxonomic diversity

Quality filtering, adaptor removal, and host read filtering were performed using shotcleaner v0.1 (42) with default parameters. For mouse fecal samples, host reads were removed by aligning to the mouse reference genome (GRCm38). A similar procedure was used for coral samples except that these reads were filtered against a concatenated version of the coral (*Acropora millepora,* Genbank: QTZP00000000.1 (43)) and symbiont (*Symbiodiniaceae* sp. clade A MAC-Cass KB8 (Tax ID: 671378, UniProt) genomes. Quality controlled sequence reads were input into HUMAnN 2.0 (44) for taxonomic and functional classification using the UniRef90 database and default parameters. HUMAnN 2.0 outputs for each community type were combined and renormalized to counts per million using HUMAnN 2.0 utility scripts before downstream analysis. High-quality reads were also taxonomically classified using Kraken2 v2.0.8-beta (45, 46) and a custom reference database that included sequences from all human, mouse (GRCm38), UniVec Core, bacteria, archaea, virus, fungi, and protozoa in the NCBI RefSeq database (accessed October 8, 2019) as well as the Symbiodinacaea sp. clade A MAC-Cass KB8 and *A. millepora* genomes.

### Statistical Analyses

Independent linear models (R::stats::lm) were used to determine how community type, library preparation method, and input DNA concentration affect variance of the resulting library characteristics including: the number of reads generated, median fragment size, library concentration, library molarity, mean read length, read duplication rate, mean read GC content, and total reads filtered and removed. Since we reasoned it was likely that interactions between the predictors exist, we employed a model selection procedure to identify the most parsimonious model for each characteristic examined. For each characteristic we built a set of models of increasing complexity: 1) a reduced model with only additive effects (eq. 1), 2) a model with interaction terms for community type, library preparation procedure, and DNA input concentration (eq. 2).

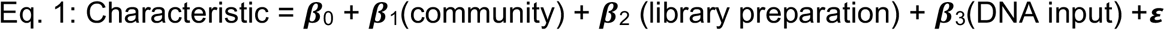

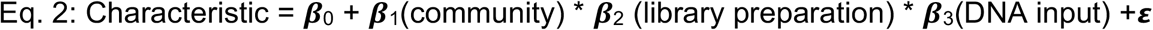

We then used Akaike information criterion (AIC) to select the most parsimonious model and analysis of variance (ANOVA) to determine the significance of each term in the selected model. Because sequencing libraries produced from distinct samples are pooled as part of the plexWell™ library preparation protocol, a single value is available for median fragment size, sequence library concentration, sequence library molarity for this method. The similarity between taxonomic profiles generated by library preparation methods and the known taxonomic composition of the ZymoBIOMICS™ Microbial Community DNA Standard was assessed by Pearson’s correlation test (R::stats::cor.test). To measure the variation in taxonomic profiles generated by different library preparation methods across soil, coral, and fecal communities, we calculated the Pearson’s correlation coefficient of generated taxonomic abundance profiles for each pair of samples. This analysis was conducted using Kraken2, a sensitive read binning tool, as MetaPhlAn2, a marker-gene-based abundance estimation tool to eliminate the possibility that mammalian biases in marker gene databases would skew results in environmental samples. We accounted for the effects of multiple correlation tests using false discovery rate (R::stats::p.adjust, method = fdr).

The additive and interactive statistical effects of library preparation and DNA input concentration on microbiome composition, as measured by the Bray-Curtis dissimilarity metric, were evaluated using PERMANOVA analysis (R::vegan::adonis, permutations = 5000, method= bray) and visualized using an ordination of principal components analysis for each community. Differences in the Bray-Curtis dissimilarity of taxonomic abundance profiles within and across library preparation methods were measured using Kruskal-Wallis tests (R::stats::kruskal.test) with a post-hoc pairwise-Wilcoxon-test (R::stats::pairwise.wilcox.test). A holm correction was used to control Wilcoxon-test family-wise error rates.

Shannon entropy and gene richness were calculated for HUMAnN 2.0 gene abundance profiles using R and vegan. Linear regression to quantified associations between gene level alpha diversity and library preparation and input DNA concentration for each community type. Associations with gene level Bray-Curtis dissimilarity, preparation method and DNA input were quantified using PERMANOVA (R::vegan::adonis, permutations = 5000, method = bray). Differences in gene abundances and metagenomic dissimilarity were quantified as with taxonomy.

## ACKNOWLEDGEMENTS

Research reported in this publication was supported by the National Science Foundation under Grant No. OCE-1933165 and BIO-2025457, as well as the National Institute of Allergy and Infectious Diseases under award number R21AI135641 and the National Institute of Environmental Health Sciences under award number R01ES030226, and a contract to the Center for Genome Research and Biocomputing from the Soil Health Institute. Library preparation kits used in this study were generously donated by PerkinElmer, Inc. and soil samples were provided by Elizabeth Rieke (Soil Health Institute).

---

**Figure S1.**
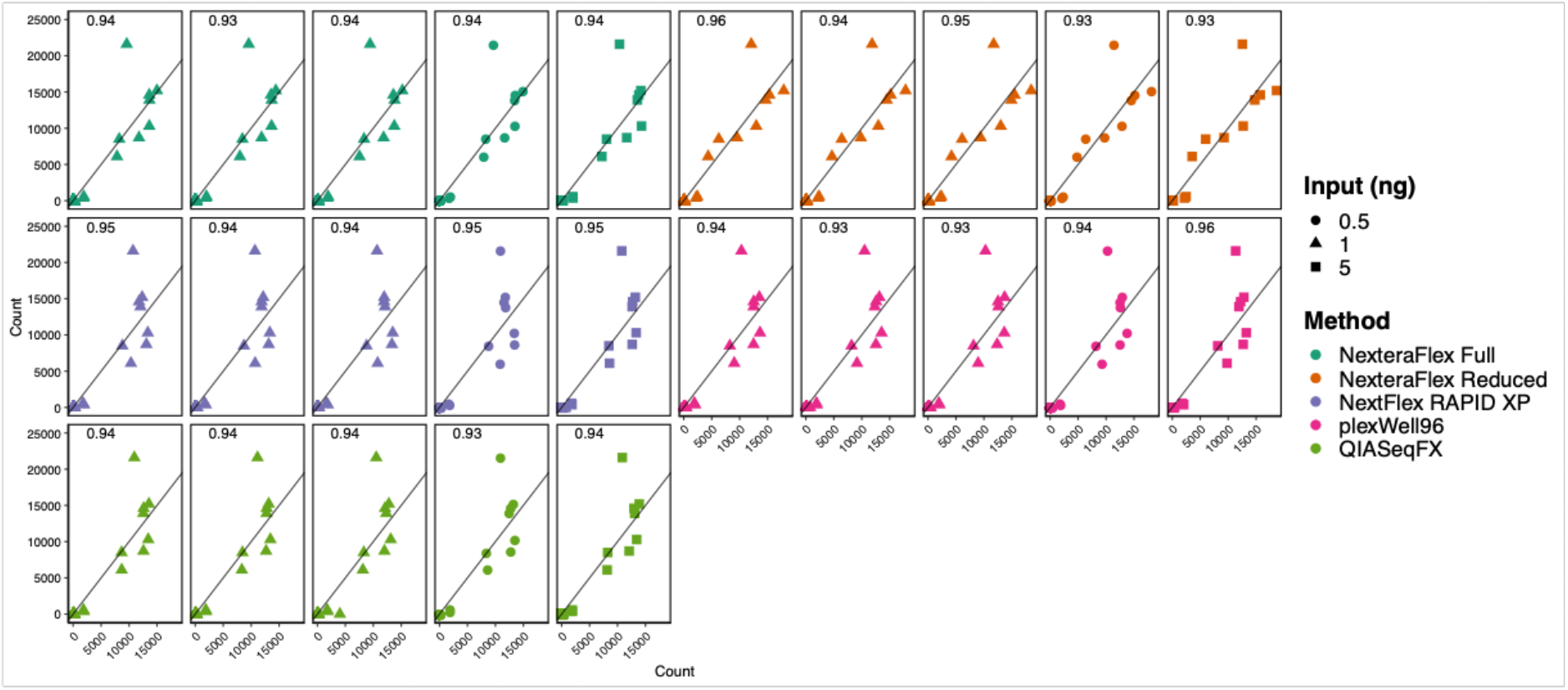
Kraken2 imputed abundance profiles strongly correlate with theoretical mock community compositions across all metagenomic library preparation methods. Scatter plots of the inferred Kraken2 taxonomic profiles and theoretical taxonomic compositions of a mock community. Library preparation methodology is indicated by point color and input concentration is denoted with point shape. Values in each panel represent the Pearson correlation coefficient.

